# DNA methylation differences between the female and male X chromosomes in human brain

**DOI:** 10.1101/2024.04.16.589778

**Authors:** Robert Morgan, Eddie Loh, Devika Singh, Isabel Mendizabal, Soojin V. Yi

## Abstract

The mechanisms of X chromosome inactivation suggest fundamental epigenetic differences between the female and male X chromosomes. However, DNA methylation studies often exclude the X chromosomes. In addition, many previous studies relied on techniques that examine non-randomly selected subsets of positions such as array-based methods, rather than assessing the whole X chromosome. Consequently, our understanding of X chromosome DNA methylation lags behind that of autosomes. Here we addressed this gap of knowledge by studying X chromosome DNA methylation using 89 whole genome bisulfite sequencing (WGBS) maps from neurons and oligodendrocytes. Using this unbiased and comprehensive data, we show that DNA methylation of the female X chromosomes is globally reduced (hypomethylated) across the entire chromosome compared to the male X chromosomes and autosomes. On the other hand, the majority of X-linked promoters were more highly methylated (hypermethylated) in females compared to males, consistent with the role of DNA methylation in X chromosome inactivation and dosage compensation. Remarkably, hypermethylation of female X promoters was limited to a group of previously lowly methylated promoters. The other group of highly methylated promoters were both hyper- and hypo-methylated in females with no obvious association with gene expression. Therefore, X chromosome inactivation by DNA methylation was exclusive to a subset of promoters with distinctive epigenetic feature. Apart from this group of promoters, differentially methylated regions in the female and male X chromosomes were dominated by female hypomethylation. Our study furthers the understanding of X-chromosome dosage regulation by DNA methylation on the chromosomal level as well as on individual gene level.

## Introduction

The presence of heterogametic sex chromosomes can cause dosage imbalance between sexes (1). For example, if all human X-linked genes were expressed equally, females would have two times the RNA products of the X-linked genes compared to males. Lyon (2) proposed that the X chromosome inactivation (XCI) evolved to silence one of the two female X chromosomes to balance the gene dosage. In mammals, XCI occurs early in female development to produce one active X chromosome (Xa) and one inactive X chromosome (Xi), via coordinated epigenetic processes (3, 4).

One of the key mechanisms in this process is DNA methylation, which refers to the addition of a methyl group (-CH_3_) to a cytosine. While the diverse roles of DNA methylation in animal genomes continue to be discovered (5-7), currently the best understood mechanism of DNA methylation is related to its role in transcriptional silencing. Specifically, increased DNA methylation of regulatory regions such as promoters and enhancers is associated with down-regulation of gene expression (8, 9). DNA methylation may be implicated in XCI through several independent mechanisms. DNA methylation may directly silence genes on the Xi, or aid the maintenance of the inactivated state (3, 4). According to this model, Xi should be more heavily methylated than the Xa. Consequently, when compared between the sexes, X chromosome DNA methylation may be elevated in the females (Xa + Xi) compared to the male (Xa). Another impact of DNA methylation is related to X-linked genes that are known to ‘escape’ XCI and express more highly in females compared to males (10-12), which may be aided by hypomethylation of the associated Xi promoters (11, 13). DNA methylation is also a key factor of epigenetic silencing of the *Xist* gene in the active X chromosome (e.g., (14, 15)).

Studies of X chromosome DNA methylation have revealed patterns that were sometimes considered contradictory. While early studies of X chromosome DNA methylation reported findings consistent with hypermethylation of Xi, a study of DNA methylation using an oligonucleotide array and using females with only one X chromosome have revealed that not all regions followed this pattern (16). Specifically, both hypermethylated and hypomethylated CpG islands were found on Xi (16). Another key study found hypermethylation of promoters and hypomethylation of gene bodies on Xi (17). It is notable that gene body DNA methylation is related to active transcription (18). The Hellman & Chess study (17) indicates that gene body hypomethylation may mark inactive genes. More recently, analyses of whole genome bisulfite sequencing data (WGBS) showed that the human, mouse, and koala female X chromosomes were globally hypomethylated compared to the male X chromosome, and that this pattern was more pronounced for gene bodies and intergenic regions (19-21). Given that the overall methylation level of the female X chromosomes is the sum of the Xa and Xi methylation and that the methylation level of the male X chromosome is that of the Xa, this pattern suggests reduced methylation of the inactive X chromosome (Xi). Therefore, the DNA methylation difference between the female and male X chromosomes is still incompletely understood. In fact, studies of DNA methylation often exclude the X chromosome, precisely due to the lack of knowledge on the impact of epigenetic patterns associated with the XCI (e.g., (22)).

This glaring gap of knowledge impedes our understanding of X chromosome regulation, which is unfortunate given the increasingly appreciated impacts of the X chromosome in health and diseases (23-26). Furthermore, variation of XCI between individuals is related to disease prevalence including cancers (27, 28). A critical limitation of several previous analyses of X chromosome DNA methylation is their bias towards subsets of CpG sites, such as those in gene promoters or CpG islands. This bias extends from array methods. As an example, the Illumina 450K array measures DNA methylation at around 500,000 CpGs, less than 2% of total CpGs in the human genome, and near promoters of genes (29, 30). However, CpG sites implicated with diseases are not necessarily found in promoters; they are often found in regions traditionally referred to as ‘non-coding’ regions of the genome (e.g., (31-33)).

Technical advances in the recent decade permit unbiased sampling of CpGs using genomic sequencing. Whole-genome bisulfite sequencing (WGBS) measures DNA methylation at single base resolution across an entire genome, thus analyzing far higher percentages of sites at genic, intergenic, and distal regions than microarray data (34). Here we analyzed previously generated 89 whole genome methylomes with an average 95% coverage of the almost 1.3 million CpGs on the human X chromosome (35). We further identified differential DNA methylation between the male and female X chromosomes utilizing this high-resolution dataset. Our analyses clearly depict both global chromosome-level and gene- and region-specific differential DNA methylation between the female and male sex chromosomes, furthering our understanding of epigenetic regulation of X chromosome inactivation.

## Results

### Global hypomethylation of the X chromosome in females associates with XCI

We used the WGBS data generated from cell populations separated using the Fluorescence activated nuclei sorting (35). Specifically, these data are from postmortem tissue dissected from BA46 of the dorsolateral prefrontal cortex, using NeuN and OLIG2 antibodies (Supplementary Table 1) representing neurons and oligodendrocytes (35). Within each cell type, neurons (NeuN+) and oligodendrocytes (OLIG2+), samples were considered by sex to form a total of four groups (NeuN+/Female, *n* = 18; NeuN+/Male, *n =* 31; OLIG2+/Female, *n* = 18; OLIG2+/Male, *n* = 22). A principal component analysis of the X chromosome DNA methylation indicates a clear separation of the male and female X chromosomes (Figure 1A). The difference between males and females explained over 70% of variation in DNA methylation in the X chromosomes. The two cell types were also clearly separated by the second principal component which accounted for 17% of variation of DNA methylation (Figure 1A). The diagnosis of schizophrenia had little effect on the variance in DNA methylation (Figure 1B), as previously shown in the analysis of autosomes (35).

**Figure 1.**
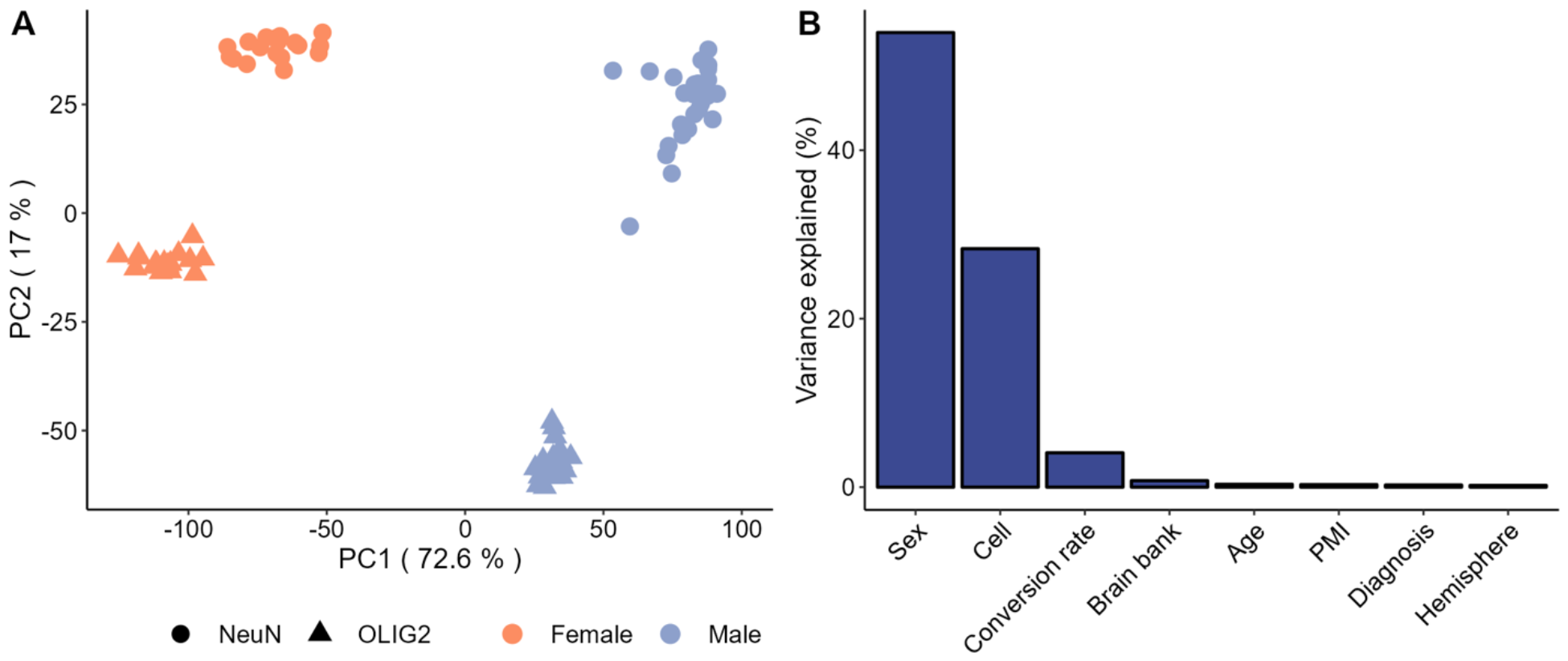
Variation of DNA methylation in the human X chromosome. (A) Principal component analysis (PCA) plots demonstrate the impacts of sex (PC1) and cell-type (PC2). (B) Variance explained by different biological variables indicate the sex and cell-type as two major factors.

Chromosome-wide (Figure 2), NeuN+ cells were more highly methylated than OLIG2+ cells, consistent with the pattern observed in autosomes (35). We observed significantly lower levels of DNA methylation of the female X chromosome compared to the male X chromosome, for both cell types (*P* < 10^−15^ for both cell types, Mann-Whitney U test, Figure 2). Comparing DNA methylation levels of autosomes and the X chromosome, it was clear that this pattern was due to the reduced DNA methylation of the female X chromosome as opposed to the increased DNA methylation of the male X chromosome (Figure 2). Consequently, we refer to this pattern as the “female X hypomethylation”. Note that female hypomethylation was not observed for the pseudoautosomal regions (PAR1 and PAR2), which recombine in both male and female germlines and do not undergo X-chromosome inactivation (36). This observation affirms that female X hypomethylation is associated with the XCI.

**Figure 2.**
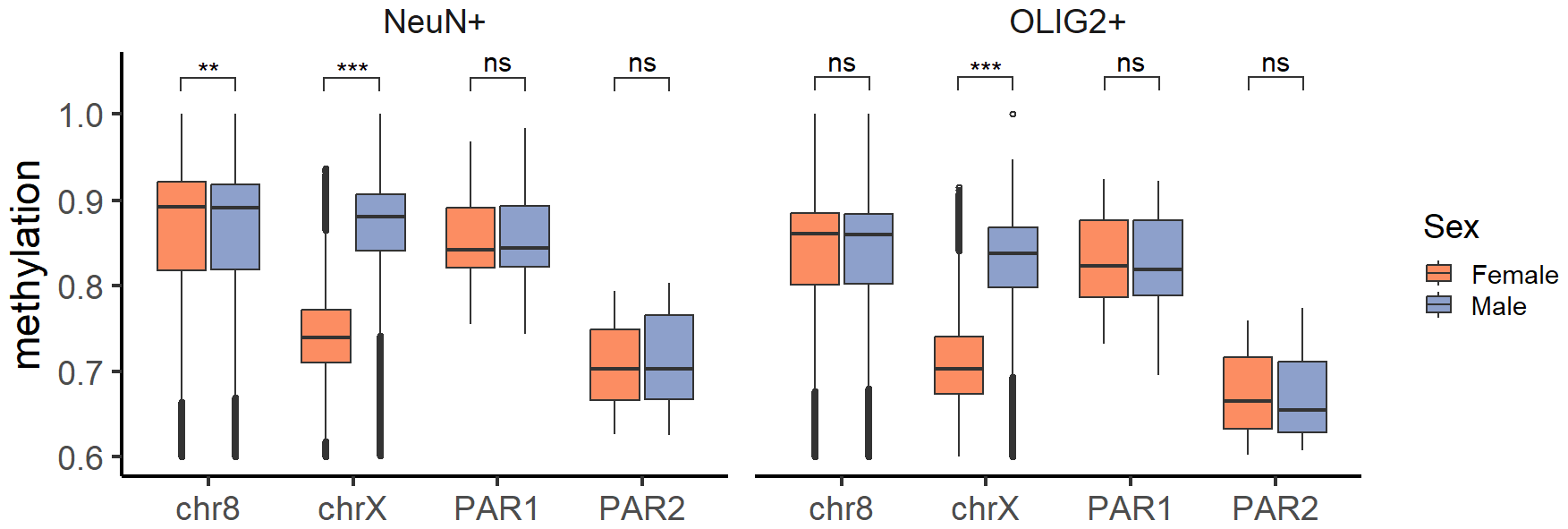
Hypomethylation of female X chromosomes in neurons (NeuN+) and oligodendrocytes (OLIG2+). Mean fractional methylation of 100kb bins across autosomes (represented by chr8 which has lengths and GC percentages close to chrX), chrX (excluding PAR regions), PAR1, and PAR2 are depicted for A) NeuN+ and B) OLIG+2 samples. Significance level * indicates *P < 0*.*05*, ** indicates *P < 1e-6*, *** indicates *P < 2*.*2e-16*, ns indicates not significant (Mann-Whitney U test).

### Differential DNA methylation and differential expression of X-linked genes

To investigate DNA methylation differences between the female and male X chromosomes in depth, we identified promoters and gene bodies of the X-linked genes. We used protein-coding genes in this analysis to avoid spuriously annotated non-coding genes. An exception to this was the *Xist* lncRNA which was included in downstream analysis due to its extremely well characterized and key role in XCI.

In contrast to the global pattern of female X hypomethylation, we observed a significant excess of female hyper-methylated promoters (447 vs. 293 female hyper-vs. hypo-methylated promoters, *P* < 10^−7^, Chi-square test, Table S2). However, it is notable that a substantial number of promoters were female-hypomethylated. On the other hand, the majority of gene bodies were female-hypomethylated (177 vs 625 female hyper-vs hypo-methylated genes, *P* < 10^−16^, Chi-square test, Table S3), consistent with the global pattern. These patterns were observed for both neurons and oligodendrocytes (Table S2, Table S3, Figure S1).

We analyzed sample-matched RNA-seq data to investigate the association between DNA methylation and gene expression of the X chromosomes (WGBS and RNA-seq data were mapped to the X chromosomes, Materials and Methods). We identified only 66 significantly differentially expressed genes out of a total 797 genes between the female and male X chromosomes (Supplementary Table 4), which affirms the active XCI and dosage compensation. In the following analyses, we examined expression differences regardless of statistical significance to understand the relationship between differential DNA methylation and gene expression. Given its direct role in silencing gene expression, we analyzed DNA methylation of promoters in the following sections.

We observed that for the majority of genes with female hyper-methylated promoters, gene expression was lower in females compared to in males, following the canonical model of transcriptional silencing by promoter methylation. This trend was significant regardless of the cell-type (308/(308+138), *P* < 10^−15^ pooled data; 280/(280+171), *P* < 10^−6^ in neurons; 303/(303+147), *P* < 10^−12^, in oligodendrocytes, Chi-square tests). However, the degree of differential DNA methylation between the female and male X chromosomes was not correlated with the degree of differential gene expression (Figure 3A, Pearson’s correlation test, *P* > 0.5), indicating that silencing of X chromosome genes via DNA methylation of promoters was not necessarily dose-dependent. In addition, while female hyper-methylated promoters were preferentially associated with female down-regulated genes, female hypo-methylated promoters were associated with both female down- and up-regulated genes (Figure 3A).

**Figure 3.**
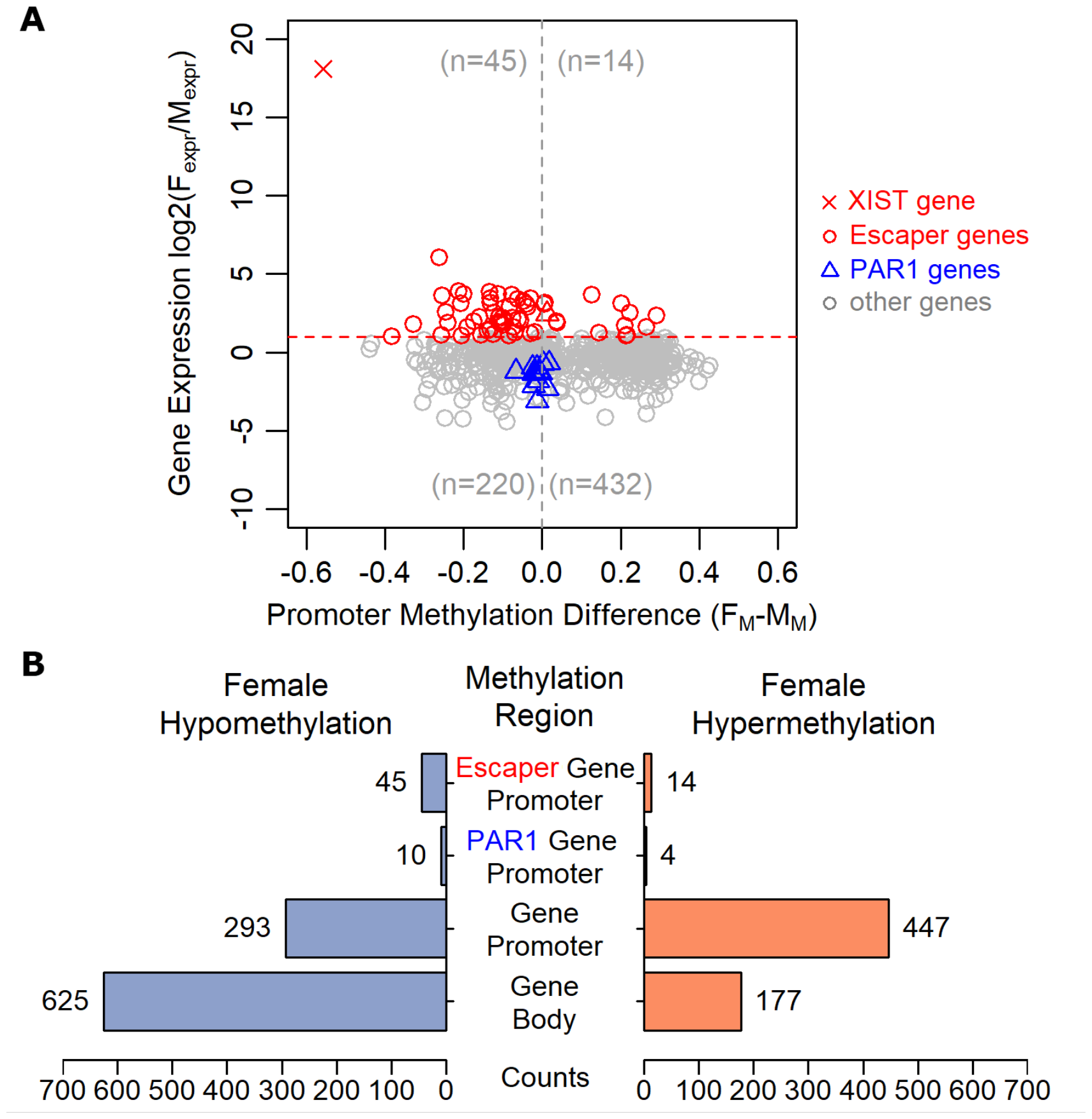
DNA methylation and gene expression differences between the female and male X chromosome linked genes. A) Scatter plot of gene expression log_2_(fold-change) (on the Y-axis) against gene promoter methylation difference on the X-axis (F_expr_ = female expression; M_expr_ = male expression; F_M_ = female methylation; M_M_ = male methylation). Each point represents a protein-coding gene. Red X indicates the *Xist* lncRNA, which is highly male-hypermethylated and female over-expressed. Red dashed line at log_2_(fold-change) = 1, indicates that genes above this line (colored in red) are more highly expressed in females than in males, which we refer to as ‘escapers’. Triangle-shaped points in blue are PAR1-linked genes, which are all more highly expressed in males compared to males, consistent with a previous study (37). B) Contrasting numbers of female hypo-vs. hyper-methylation regions.

### Differential DNA methylation between female and male X chromosomes implicated in escapers but not in the down-regulation of PAR genes

Some X-linked genes are known to ‘escape’ X chromosome inactivation, leading to increased female expression (11, 12). In our data, there were 59 genes whose expression was greater than two-fold higher in females compared to males (note that only 5 out of these 59 were statistically significant at *P* < 0.05). Remarkably, the majority of these escapers (45 out of 59, 76.3%) were female hypo-methylated (Figure 3A, 3B). This observation is consistent with the idea that hypo-methylation of promoters could contribute to escapers.

Genes located on the pseudoautosomal region 1 (PAR1) of the X are transcriptionally down-regulated in females compared to males, potentially due to the XCI mechanism extending to PAR1 (37). Consistent with this previous study, we observed female drown-regulation of PAR1 genes in both cell types (Figure 3A). If DNA methylation was involved in this process, we expect PAR1 gene promoters hypermethylated in females than in males. In contrast, promoters of PAR1-linked genes were predominantly hypomethylated in females compared to males, although the extent of hypomethylation was not as pronounced as in the rest of X chromosome genes (Figure 3A). These results indicate that the down-regulation of PAR1-linked genes of female X chromosomes is not driven by DNA methylation (Figure 3A, B). Data from the two cell types provided similar results when analyzed combined (as in Figure 3) or separately (Supplementary Figure 2).

### Differentially methylated regions across cell types and sex

Above we have investigated differential DNA methylation between females and males using well-annotated promoters and gene bodies. WGBS data also provides an opportunity to examine nearly every CpG on the X chromosome. Utilizing this extensive data, here we identified differentially methylated regions (DMRs) across the whole X chromosome (Materials and Methods). Following stringent filtering steps and correcting for multiple testing, we obtained a total of 16,183 DMRs that are differentiated between the female and male X chromosomes (referred to as ‘sex-DMRs’), 3,287 DMRs between the two cell types (referred to as ‘cell type-DMRs’), and 40 DMRs that indicated interactions between sex and cell-type (Materials and Methods). Given the small number of the interaction DMRs, in the next sections we focus on sex- and cell type-DMRs. Regardless, genomic locations of DMRs and their annotations, including the associated genes, are shown in Supplementary Table 5.

While many of these DMRs are found near genes (promoters, exons and introns), between 43-37% of those were found in intergenic regions (Figure 4A, B). We examined the direction of differential DNA methylation of these DMRs. Sex-DMRs were dominated by female hypomethylation (Figure 4C), which is consistent with the global hypomethylation of the female X compared to the male X (Figure 1). Nevertheless, among the sex-DMRs, those that occur in promoters include a large number of female hyper-methylation (Figure 4C). Over 80% of X-linked genes (647 out of 791) harbored sex-DMRs, consistent with the broad, chromosome-wide differential DNA methylation of the female and male X chromosomes contributing to XCI.

**Figure 4.**
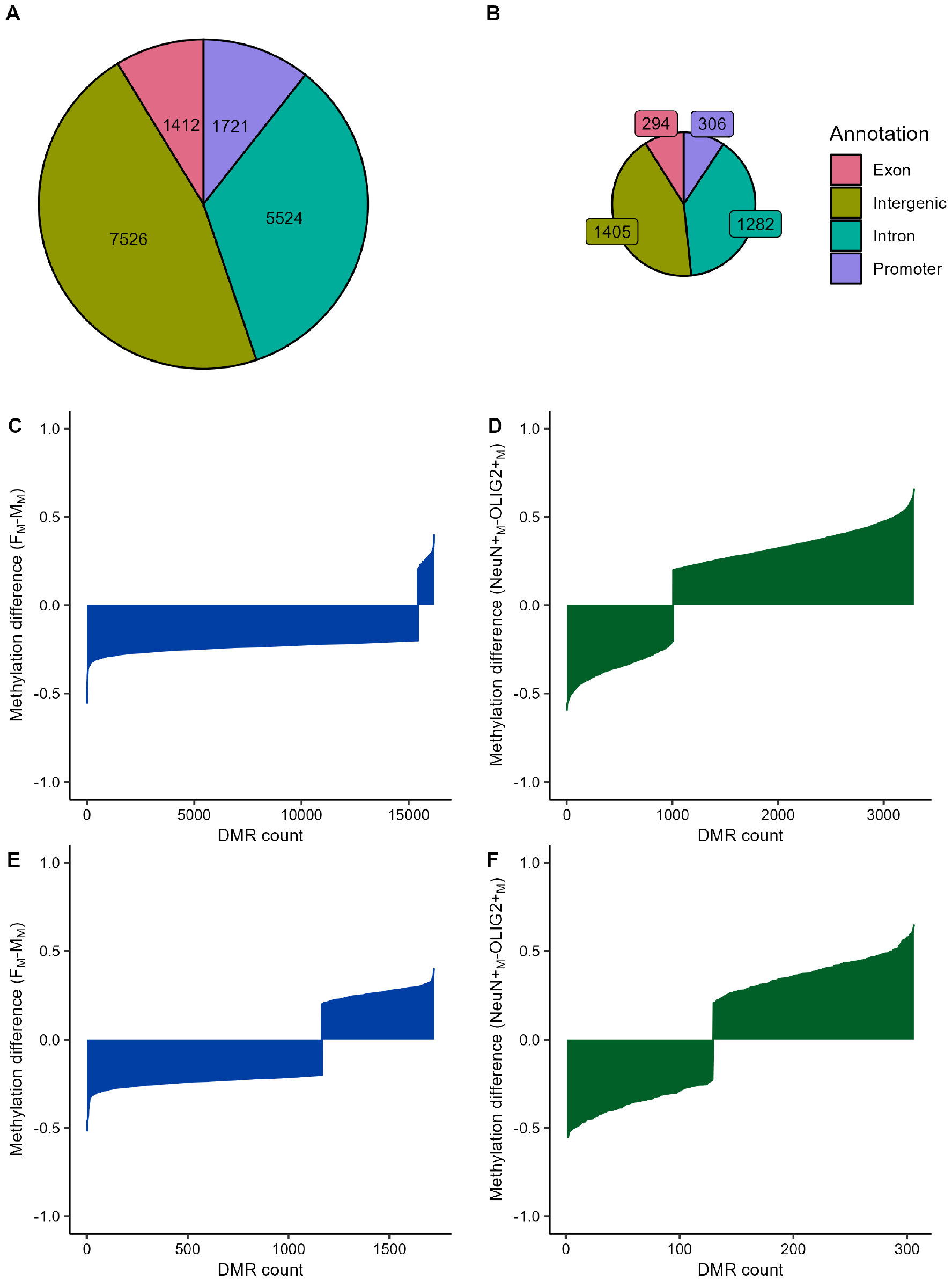
Differentially methylated regions (DMRs) between the female and male X chromosomes. (A) The distribution of sex-DMRs and (B) cell-type DMRs across functional regions. (C) The majority of sex-DMRs are hypomethylated in females (F_M_ = female methylation; M_M_ = male methylation). (D) The majority of cell-type DMRs were more highly methylated in neurons compared to oligodendrocytes (NeuN+_M_ = neuron methylation; OLIG2+_M_ = oligodendrocyte methylation). When we examined promoter DMRs, (E) a large number of promoter DMRs were hypermethylated in females compared to males. (F) cell-type promoter DMRs were slightly dominated by neuron hypermethylation.

In contrast, cell-type DMRs were predominantly biased toward hypermethylation in neurons (Figure 4C). A total of 452 X-linked genes harbored at least one cell-type DMRs, among which 232 genes significantly differentially expressed (referred to as ‘DE’ hereafter) between the two cell types (Supplementary Table 6). This represents a highly significant excess of cell-type DMRs for DE genes (Table 1). DE genes also had a greater number of DMRs (2.50 DMR/gene for DE genes vs. 1.34 DMRs/gene for non-DE genes), which was highly significantly different from a bootstrapping test (Z-score of 7.32 or *P* <10^−10^; Materials and Methods). Genes associated with strong (methylation difference >0.3) promoter cell-type DMRs were enriched in gene ontologies including *anatomical structure development, developmental process*, and *cellular component organization*, and *neuron development* (FDR < 0.05, Fisher’s Exact Test). Our analysis provides a novel list of X-linked differentially expressed genes between neurons and oligodendrocytes (Supplementary Table 6) and attests to the strong roles of DNA methylation in regulating cell-type specific gene expression (35).

**Table 1.** Genes differentially expressed between neurons and oligodendrocytes (‘DE genes’) are associated with differentially methylated regions (*P* < 10^−7^ by Fisher’s exact test).

|  | DE gene | Non-DE gene |
| --- | --- | --- |
| Has associated DMR | 232 | 220 |
| No associated DMR | 106 | 233 |

### Female hypermethylation is driven by lowly methylated promoters

Above we have demonstrated that promoters on the female X chromosome undergo both hyper- and hypo-methylation. In this section, we show that whether the promoters undergo hyper- and hypo-methylation is not random. Rather, the innate functional bimodality of promoters associated with the direction of DNA methylation on the female X chromosome, with distinct consequences in associated gene expression.

Specifically, previous studies have shown that promoters in the human genome consist of two distinctive groups classified by their DNA methylation levels (38, 39). Briefly, one group is devoid of DNA methylation (we will refer to these “lowly methylated promoters” or LMPs) (Figure 5A, Supplementary Figure 3). The other group of promoters are generally highly methylated (here referred to as “highly methylated promoters” (HMPs)) (Figure 5A, Supplementary Figure 3). These two groups of promoters are known to exhibit different genomic and functional characteristics, such that LMPs are typically broadly expressed across tissues with housekeeping functions, while HMPs tend to function tissue-specifically (38, 39). Consistent with these previous studies, X-chromosome promoters in our data were clearly distinguished as low and high levels of DNA methylation (Figure 5A, Supp Figure 3).

**Figure 5.**
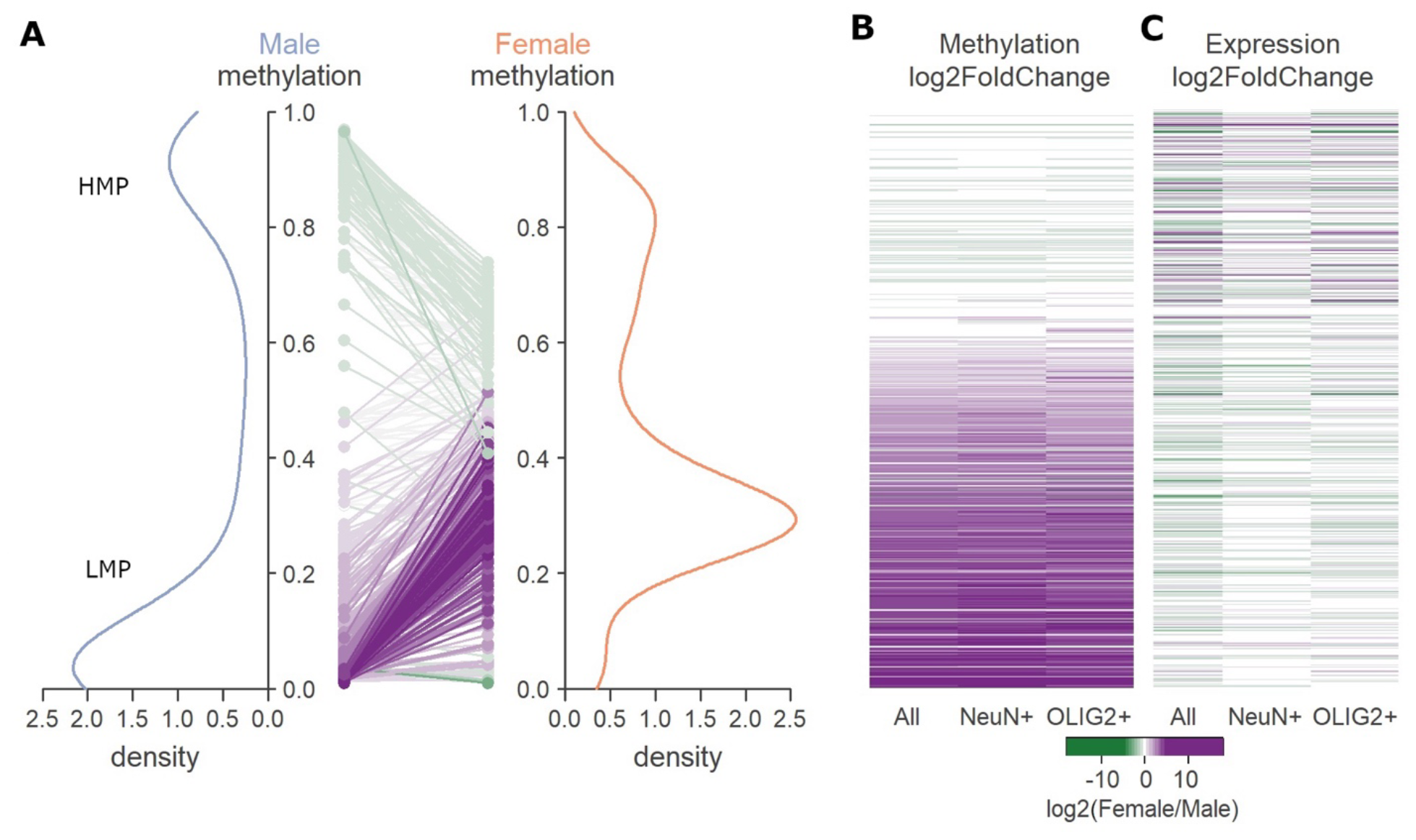
(A) Promoters of human genes can be divided into lowly (LMPs) and highly methylated (HMPs) subsets in both males and females (also see Supplementary Figure 3). Only LMPs undergo female hypermethylation, as indicated by male to female changes as colored lines (dominated by purple: increase of methylation from male to female X; using the same color scale as B and C). HMPs on the other hand are female hypomethylated (dominated by green: decrease of methylation from male to female; using the same color scale as B and C). (B) Methylation changes for each promoters, where green is female hypomethylation and purple is female hypermethylation. Promoters are sorted in the same order as in A, thus lowly methylated promoters are on the bottom and highly methylated promoters are near the top. The first column is for neurons and oligodendrocytes combined, followed by neuron and oligodendrocyte specific patterns. (C) Expression change for corresponding gene as in A and B demonstrates that the female hypermethylated promoters show little gene expression difference, as expected from the XCI. On the other hand, female hypomethylated promoters are linked to both up and down regulated genes.

Remarkably, we observed that female X hyper-methylation occurred nearly exclusively for LMPs (Figure 5A). This pattern was consistent in neurons and oligodendrocytes (Figure 5B). HMPs almost always showed female X hypomethylation (Figure 5A, 5B). Integrating with gene expression, only LMPs were associated with balanced expression between the male and female X (Figure 5C). In contrast, HMPs were associated with both up- and down-regulated genes (Figure 5C). Therefore, XCI by DNA methylation was specific to the lowly methylated promoters.

## Discussion

In this study, we addressed the question of differential DNA methylation between the female and male X chromosomes. Due to the XCI, the methylation measured from the female X chromosome corresponds to the combined signatures on the Xa and Xi, while the male X chromosome represents the information from the Xa alone. Using the extensive, nucleotide-resolution DNA methylation data from neurons and oligodendrocytes, here we report the following findings.

First, we demonstrate that the female X chromosome is globally hypomethylated compared to the male X chromosome. Synthesizing our observation with previous results, we propose that this pattern reflects hypomethylation of gene bodies and intergenic regions of the inactive X chromosome (Xi) and that it is a consistent key feature of XCI. Hellman and Chess (17) demonstrated Xi was hypomethylated in gene bodies using arrays. In another study, Cotton et al. (40) also reported female hypomethylation of non-CpG island probes in placenta, using an array of 1505 CpGs. Cotton et al. (41) further observed hypomethylation of Xi for dosage-compensated genes, using the Illumina 450k array. Recently, a study of allele-specific methylation in a specific mouse line with nonrandom XCI showed that intergenic regions of the Xi tended to be hypomethylated (21). Our results, using unbiased, chromosome-wide approach, reveals that the entire female X chromosome is hypomethylated, indicating that the aforementioned previous results reflected a major feature of sex difference in the X chromosome DNA methylation. We propose that this is due to a potentially general (but not exclusive) association between transcriptional activity and DNA methylation outside of promoters and CpG islands. Gene bodies of highly expressed genes are heavily methylated (8, 18, 42), including those on the X chromosome that escape XCI (41). Heavy DNA methylation is generally associated with the suppression of spurious transcription within gene bodies (42) and of transposable elements (43), which are prevalent in intergenic regions (44). Therefore, heavy DNA methylation may be a feature of transcriptionally active chromosomal regions, although the exact nature of the causative mechanism remains to be resolved in future studies. Transcriptionally less active regions such as the Xi may more predominantly rely on other epigenetic factors rather than DNA methylation to maintain their genome integrity. Consistent with this idea, it was recently shown that the Y chromosome, which harbors only a small number of genes and thus transcriptionally inactive across the majority of its length, exhibits significantly reduced DNA methylation compared to the X chromosome and pseudoautosomal regions (45). We caution however that there likely exist gene by gene exceptions; for example, some extremely transcriptionally active short genes are devoid of DNA methylation (46).

Second, we show that promoter DNA methylation, a main component of XCI, is limited to a subset of genes. Specifically, female hypermethylation and the subsequent dosage balance between the female and male X was nearly exclusive to genes harboring lowly methylated promoters. The other group of highly methylated promoters were associated with both up- and down-regulation of X-linked genes, including escapers. Therefore, while XCI by DNA methylation is prevalent, it is not a universal mechanism on the female X. Our findings can consolidate, and offer clear explanations for some previous studies. Sharp et al. (16) used an oligonucleotide array to show that while a large number of CpG islands were hypermethylated in Xi, some CpG islands were hypomethylated in Xi. Given that CpG islands are generally found in lowly methylated promoters (38, 39), this observation is consistent with our proposal. Using the 450K array, Cotton et al. (41) showed that promoters with high CpG density (which tend to be HMPs) were intermediately methylated in female X while lowly methylated in male X. While these previous studies used microarray methods to examine pre-determined CpG positions, we examined nearly all CpGs on the X chromosome and demonstrated that female hypermethylation is highly confined to lowly methylated subset of promoters (Figure 5).

Our study utilized unbiased, chromosome-wide DNA methylation maps and matched RNA-seq data to uncover new patterns and reconcile previous observations which were based on small scale arrays. We also observed clear cell-type differences, amid the above general patterns in neurons and oligodendrocytes (Figure 2). In addition to the neuron hypermethylation as was observed in autosomes (35), we found a large number of differentially methylated regions that were preferentially associated with differential gene expression (Figure 4, Table 1). When we compared our lists of escapers and differentially expressed genes in neurons and oligodendrocytes with those from other cell types, few genes were overlapped (Supplementary Figure 4 and 5). Therefore, it is pertinent that unbiased investigation of DNA methylation and gene expression across broad cell types is needed to mechanistically understand how XCI is regulated by DNA methylation in different cell context.

## Materials and Methods

### WGBS data

Raw reads were downloaded from the link provided by (35). These are methylomes sequenced from DNA samples collected postmortem from Brodmann area 46 (BA46) of the dorsolateral prefrontal cortex. Methylome libraries were constructed in 150 bp paired end reads and sequenced on Illumina HiSeq 2500 and HiSeqX. We selected 89 methylomes with >95% bisulfite conversion rates for further analyses (Supplementary Table 1).

### Whole-genome bisulfite sequencing data processing

Adapters and low-quality reads were removed from reads with TrimGalore v0.4.1 (The Babraham Institute) using default parameters. To remove the spike-in control, reads were mapped to the PhiX genome using Bismark v0.14.5 (47) and bowtie v2.4.4 (48).

### Mapping WGBS data to sex chromosomes

To map the WGBS data to sex chromosomes, PAR1 and PAR2 were hard-masked on chrY so reads were only mapped to the chrX PAR regions. Reads were then mapped to the GRCh38 (build 38.p12) human reference genome with Bismark v0.23.1 and bowtie v2.4.4 (48).

Alignments that share start and end positions were deduplicated with Bismark and SAMtools v1.9 (49). Methylation extraction was performed on the deduplicated reads for each possible C context (CpG, CHG, and CHH) with Bismark. From this extraction, a genome-wide cytosine report was generated. Statistical analysis on cytosine reports were performed with the bioconductor library bsseq (50).

### Filtering WGBS data

From the 1000 Genomes phase 1 SNPs (The 1000 Genomes Project Consortium, 2012), we identified C>T and G>A polymorphisms (allele frequency > 1%) at 147,830 CpG loci on the X chromosome. These loci were excluded from further analysis since their polymorphisms are consistent with bisulfite conversion and could generate false positives. Loci with < 5x coverage in more than 20% of samples for at least one sample group (F/NeuN+, F/OLIG2+, M/NeuN+, M/OLIG2+) were also removed.

### Differentially methylated positions and regions

We identified DMPs and DMRs across sex (female vs. male), cell-types (NeuN+ vs. OLIG2+), and sex/cell-type interactions. We used the Bioconductor package DSS (2.46.0) which considers biological covariates and sample-level differences in read coverage (51). Covariates included in our model definition were age, gender, brain hemisphere, postmortem interval (PMI), bisulfite conversion rates, brain bank, and genetic ancestry. Age and PMI were considered as three-level categorical variables. Genetic ancestry was defined as the first 10 genetic PCs computed from WGS samples for the same individuals (35).

DMPs were identified from the remaining set of CpG loci using a Bonferroni correction on P < 0.05. We identified a total of 435,846 sex, 86,832 cell-type, and 21,601 sex-cell-type interaction differentially methylated positions (DMPs) across the X chromosome. From the DMPs, DMRs were constructed with a minimum length of 50 bps and a minimum of 5 CpGs. DMP and DMR calling was done for three sample group comparisons: (1) sex (F vs. M), (2) cell-type (NeuN+ vs. OLIG2+), and (3) sex/cell-type interaction (F/NeuN+ vs. F/OLIG2+ vs. M/NeuN+ vs. M/OLIG2+). DMRs were subject to the same Bonferroni correction on *P* < 0.05 as the DMPs. These DMPs were analyzed with DSS to construct 24,046 sex, 3,836 cell-type, and 764 sex-cell-type interaction differentially methylation regions (DMRs). We applied an additional filtering step that required each DMR to meet or exceed 20% methylation difference averaged across its CpG loci (where the difference is between Female v. Male, NeuN+ vs. OLIG2+, or, in the case of sex-cell-type interaction, both comparisons), Due to the low number of sex-cell interaction DMRs that passed our filtering, we elected to focus our further analysis on the sex and cell-type DMRs. Using the above filtering, we identified 16,183 sex, 3,287 cell-type, and 40 sex/cell-type interaction DMRs.

### Functional annotation of DMRs

We assigned DMRs a functional category based on the nearest gene. Annotations were queried by ChIPSeeker from an Ensembl v108 hg38 TxDb generated with GenomicFeatures (52, 53). Promoters were defined as the range -2000 bp to +100 bp of a gene’s TSS. Functional regions considered were distal intergenic, intron, exon, promoter, downstream, 5’ untranslated region (UTR), and 3’ UTR.

### RNA-seq Results Analysis

RNA-seq data from matching samples (35) was downloaded from the Gene Expression Omnibus (accession no. GSE108066). Raw sequencing reads were first quality checked and trimmed using Trim Galore (v0.6.7; a wrapper program implementing Cutadapt v4.2 and FastQC v0.11.9). The trimmed reads were then directly quantified using Salmon v1.10.0 under the mapping-based mode, where a Salmon index was first built using a decoy-aware transcriptome generated from the Ensembl v108 hg38 annotation (53). The transcript-level expression read counts generated by Salmon was aggregated into gene-level read counts, after which a differential gene expression analysis was performed using DESeq2 (v1.34.0)

### Bootstrapping Analysis for Preponderance of DMRs in DE genes

Among the 791 X-linked genes in the cell-type DE analysis, 338 were significant at FDR<0.05, while the remaining 453 were not significant. Among the cell-type DMRs, 844 were annotated to the 338 DE genes (or mean of 2.50 DMRs/gene), and 608 were annotated to the 453 non-DE genes (or mean of 1.34 DMRs/gene). A 100,000 iteration bootstrap sampling was performed, whereby in each iteration, 338 genes were randomly sampled with replacement from the 791 X-linked genes, and the number of DMRs annotated to them were recorded. This yielded a distribution of bootstrapped DMR counts with a mean of 505.2 and standard deviation of 46.3. A Z-score of 7.32 (or p-value=2.46e-13) was therefore calculated from our observed 844 DMRs (annotated to the 338 DE genes), leading to our conclusion that DE genes do indeed have significantly more DMRs associated/annotated to them.

## Acknowledgement

We thank Dr. Hyeonsoo Jeong for technical help and critical feedback. RM initiated an earlier version of this study as a part of his undergraduate thesis at the College of Computing of Georgia Institute of Technology. This research was partly supported by NSF (EF-2021635), NIH (HG011641), and ICB (27KK01) grants to SVY.

