## Supplementary Figures for "DNA methylation differences between the female and male X chromosomes in human brain"

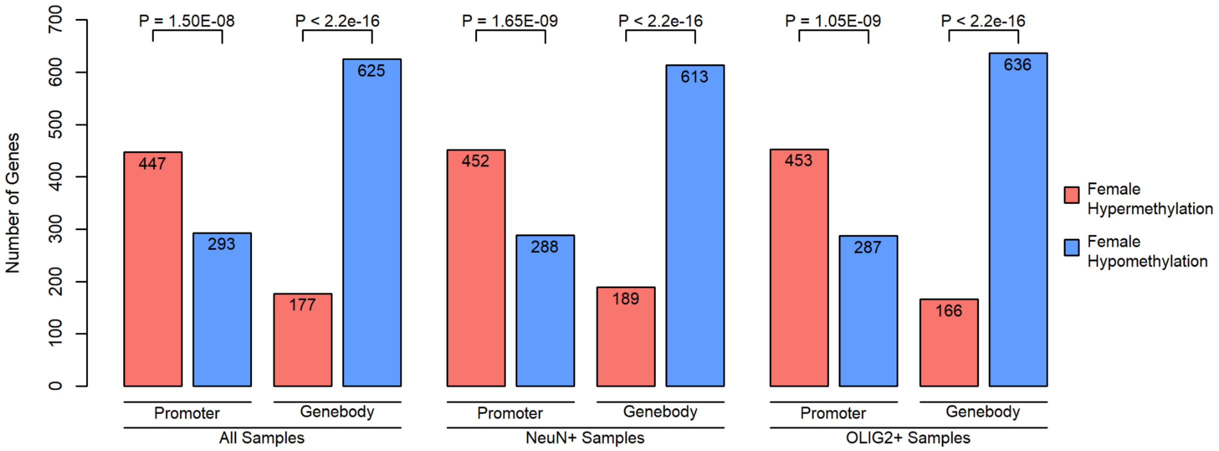


**Supplementary Figure 1.** The number of X-linked gene bodies and promoters that experience female hypermethylation and hypomethylation in all samples considered, versus those from NeuN+ and OLIG2+ samples, separately. In all cases, promoters are dominated by female X hypermethylation, while gene bodies are dominated by female hypomethylation.

**A) NeuN+ samples only**

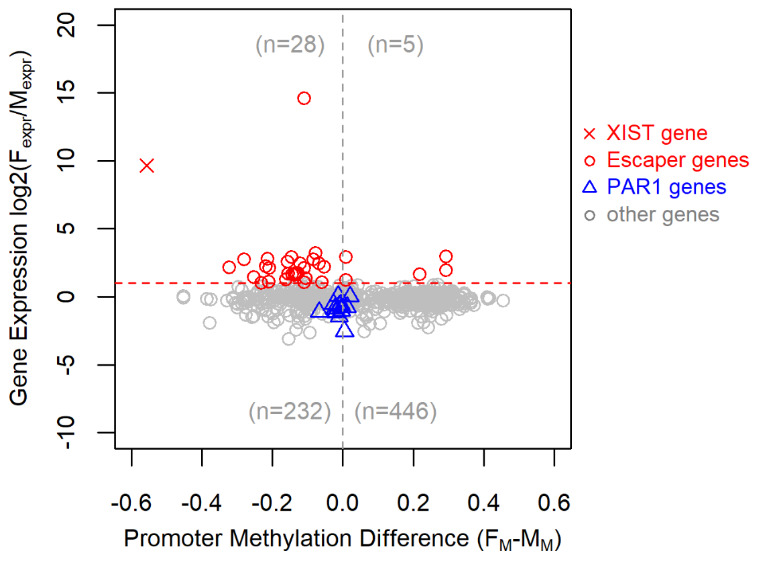


**B) OLIG2+ samples only**

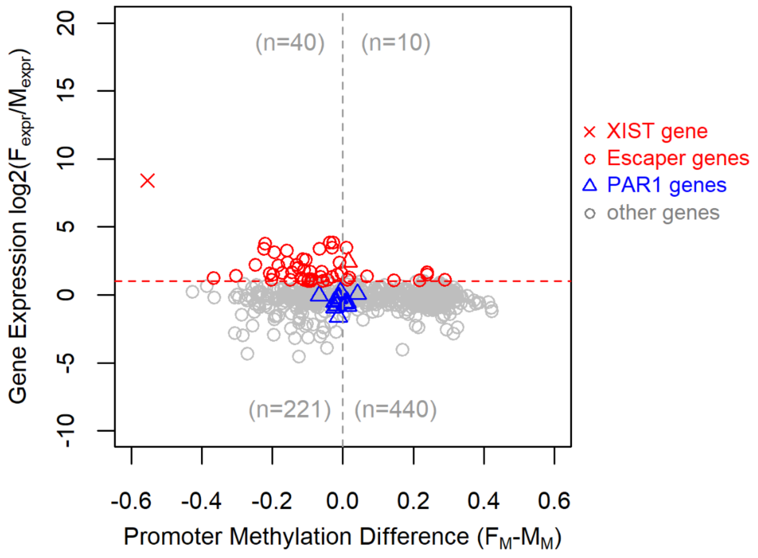


**Supplementary Figure 2.** Differential DNA methylation of male and female promoters when neurons and oligodendrocytes were examined separately. The resulting patterns are highly similar to those from pooled samples (Figure 3 in main text), and indicate that escapers are generally female hypomethylated, and PAR1 genes do not show strong differential DNA methylation.

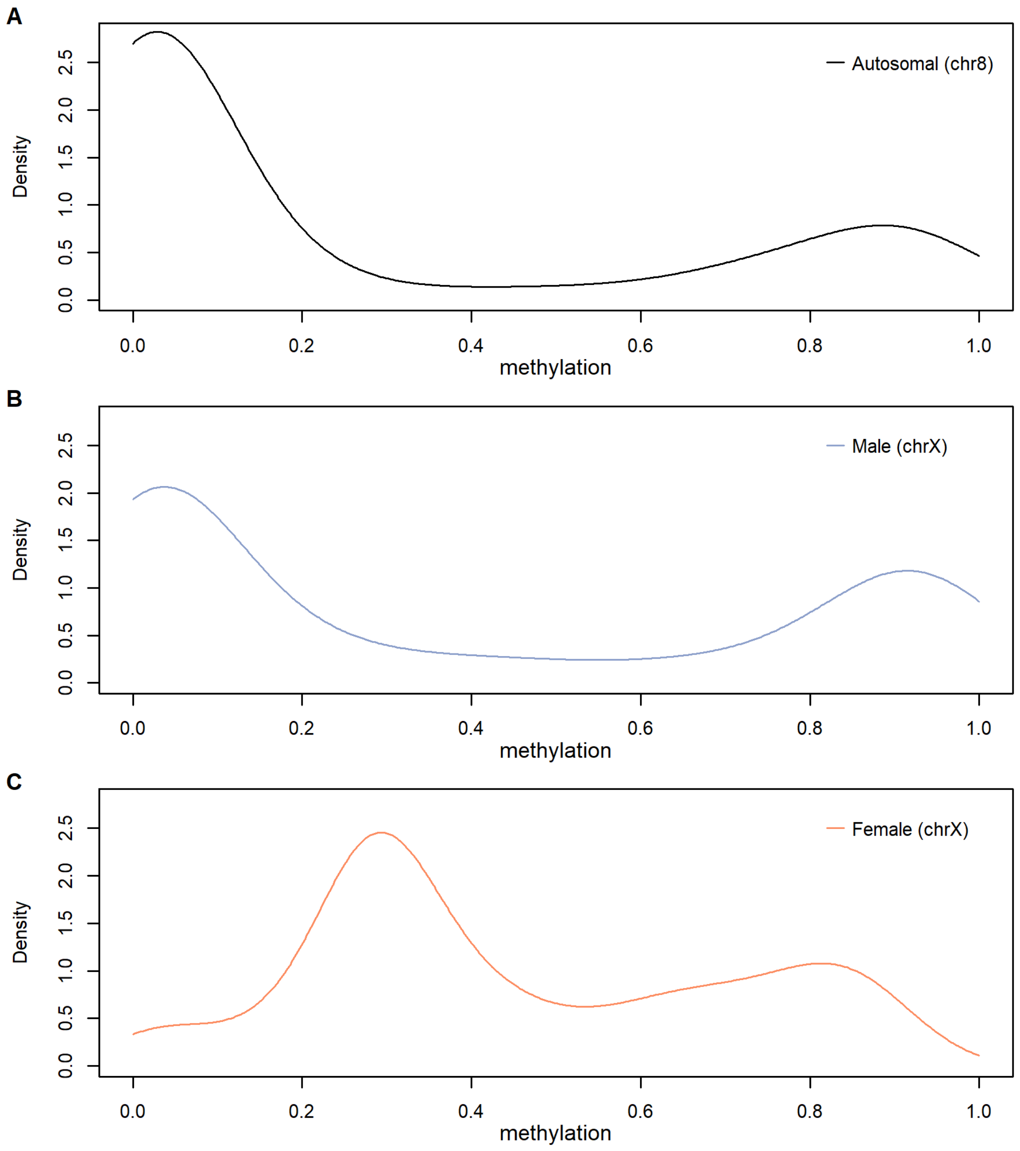


**Supplementary Figure 3.** Density plots of promoter methylation from a representative autosome (Chr 8) and from the male and female X chromosomes demonstrate clear ‘bimodality’ of lowly and highly methylated promoters. These observations are consistent with previous studies (e.g., Elango and Yi 2008, Weber et al. 2006).

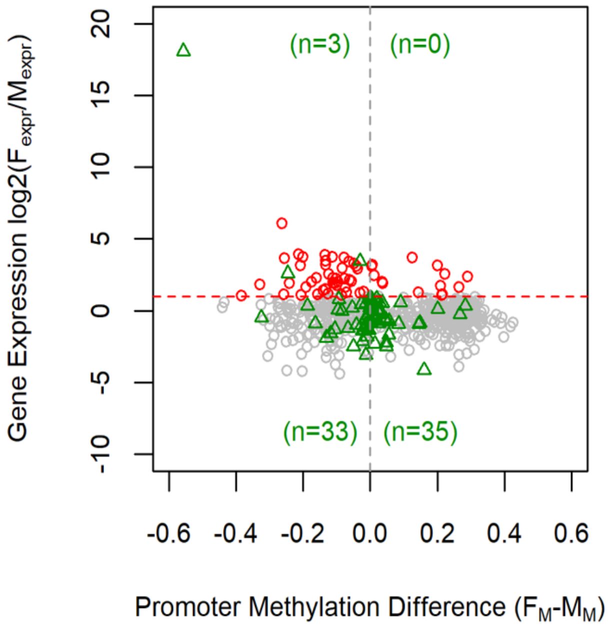


**Supplementary Figure 4.** Escaper genes identified inTukiainen et al. (2017) which are mostly from fibroblasts and lymphoblastoid cell lines are depicted in green and compared to DNA methylation difference in our data from neurons and oligodendrocytes. There was little overlap between escapers in our data set (red) and those from Tukiainen et al. (2017) (green).

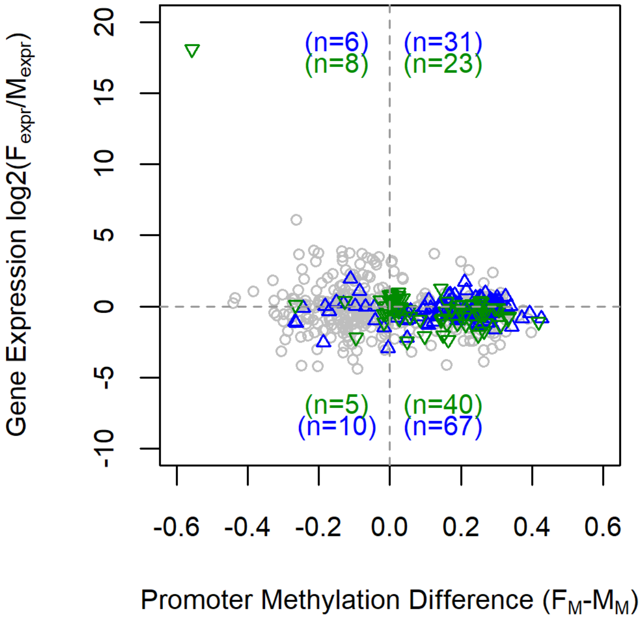


**Supplementary Figure 5.** Differentially expressed genes Wingo et al. (2023) which were identified from identified proteomics analysis, in relation to DNA methylation difference and gene expression in our matched data. Female overexpressed genes (green) and male overexpressed genes (blue) in Wingo et al. (2023) had little consistency with the distribution of DNA methylation and gene expression obtained from the BA 46 region.
